# A Detailed View of KIR Haplotype Structures and Gene Families as Provided by a New Motif-based Multiple Sequence Alignment

**DOI:** 10.1101/2020.08.07.242305

**Authors:** David Roe, Cynthia Vierra-Green, Chul-Woo Pyo, Daniel E. Geraghty, Stephen R. Spellman, Martin Maiers, Rui Kuang

## Abstract

Human chromosome 19q13.4 contains genes encoding killer-cell immunoglobulin-like receptors (KIR). Reported haplotype lengths range from 67 to 269 kilobases and contain 4 to 18 genes. The region has certain properties such as single nucleotide variation, structural variation, homology, and repetitive elements that make it hard to align accurately beyond single gene alleles. To the best of our knowledge, a multiple sequence alignment of KIR haplotypes has never been published or presented. Such an alignment would be useful to precisely define KIR haplotypes and loci, provide context for assigning alleles (especially fusion alleles) to genes, infer evolutionary history, impute alleles, interpret and predict co-expression, and generate markers. In order to extend the framework of KIR haplotype sequences in the human genome reference, 27 new sequences were generated including 24 haplotypes from 12 individuals of African American ancestry that were selected for genotypic diversity and novelty to the reference, to bring the total to 68 full length genomic KIR haplotype sequences. We leveraged these data and tools from our long-read KIR haplotype assembly algorithm to define and align KIR haplotypes at <5 kb resolution on average. We then used a standard alignment algorithm to refine that alignment down to single base resolution. This processing demonstrated that the high-level alignment recapitulates human-curated annotation of the human haplotypes as well as a chimpanzee haplotype. Further, assignments and alignments of gene alleles were consistent with their human curation in haplotype and allele databases. These results define KIR haplotypes as 14 loci containing 9 genes. The multiple sequence alignments have been applied in two software packages as probes to capture and annotate KIR haplotypes and as markers to genotype KIR from WGS.

## Introduction

KIR gene names reflect their protein structures(1). First the prefix “KIR” (killer-cell immunoglobulin-like receptors), followed by the number of extracellular domains (“2D” or “3D”), followed by a short or long intracellular domain (“S”, L”), followed by a number indexing their naming. *KIR2DS1*, for example, has two extracellular domains and a short intracellular domain; it is the first gene named with that structure, and *KIR2DS2* is the second. KIR haplotypes have no official nomenclature, although the publications that have contributed most of the haplotypes in the human genome reference use a convention set in publications by Pyo et al. in 2010(2) and 2013(3). The haplotype names reflect the two-part structure of the region: a proximal centromeric (‘c’) region is paired with (‘~’) a distal telomeric (‘t’) region. Each region is a variant of a ‘A’ haplotype or ‘B’ haplotype family, followed by a number indexing their naming. The haplotype named ‘cB02~tA01’, for example, is comprised of the second centromeric B region (“cB02”) in cis with (“~”) the first telomeric A region (“tA01”).

The Immuno Polymorphism Database for KIR (IPD-KIR) names KIR gene alleles, records their DNA-RNA-protein relationships, and annotates each gene’s alleles in a multiple sequence alignment(4). There is no equivalent for KIR haplotypes. Although there are many publications that analyze gene or intergene allele alignments, none report full haplotype MSAs, and therefore the haplotype nomenclature has not been formalized. Allele gene assignments are evaluated largely by amplification primers or sequence similarly to other alleles, as opposed to location in its haplotype. To help solve these issues, we present a simple bioinformatics approach to represent KIR haplotypes as a string of alignment-based motifs.

In our recent preprint describing a KIR long-read assembler(5), we show that 18 120 bp sequences can be used to capture full haplotype KIR DNA from PacBio circular consensus sequencing (CCS) reads. In this manuscript, we show that when those probe sequences are aligned to assembled haplotypes, the pattern of the probes (‘motifs’) provides a structural annotation. These alignment locations of the motifs allow the haplotype sequences to be represented accurately and efficiently at the structural level. We leveraged that annotation to align the 68 published human KIR haplotypes and one chimpanzee haplotype to within an average of 2398 bases, and then we micro aligned each locus for precise full haplotype multiple sequence alignment. The results are concordant with the annotation in the human genome reference and reveal 13 structural haplotypes for the 68 human haplotype sequences. The MSA also shows the haplotypes have 14 loci containing 9 genes.

The MSA includes this study’s contribution of 27 new haplotypes to the human genome project, including 24 haplotypes from 12 individuals of African American ancestry.

## Materials and Methods

### Source sequences and annotation

The source sequences consisted of two newly assembled haplotypes from an Ashkenazim individual (from the assembler preprint), along with all 66 full-sequence alternative haplotypes in the human genome reference(6)(2)(3)(7) and a chimpanzee haplotype (GenBank accession AC155174.2)(8). The two Ashkenazim haplotypes were assembled with PacBio reads obtained from the Genome In a Bottle (GIAB) consortium(9) and have not yet been submitted to genetic databases. All other haplotypes were generated by physically separating chromosomes via fosmid cloning and then sequencing and assembling fragments into full haplotype sequences. The 68 include 27 new sequences first described in this manuscript, including 24 haplotypes from 12 individuals of African American ancestry that were selected by genotypic diversity and/or lack of representation in the human genome reference. An additional 3 haplotypes (1 Asian, 2 European) of opportunity from two individuals were contributed by Scisco Genetics, who sequenced all non-GIAB haplotypes following previously reported protocols(2)(3). The GenBank entries also provided structural and allelic annotation of all the human haplotypes, except for the two Ashkenazim haplotypes. Broken down by population, the haplotype counts include 31 African or African American (AFA), 1 Asian, 2 Ashkenazim, 2 Chinese, 22 European, 2 Guarani South Amerindian, 2 Hispanic, 2 Spanish Gypsy, and 4 unknowns. The structures are depicted schematically in Figure 1, excluding cA01~tB04, which contains a large insertion and was excluded for compact visualization.

**Figure 1.**
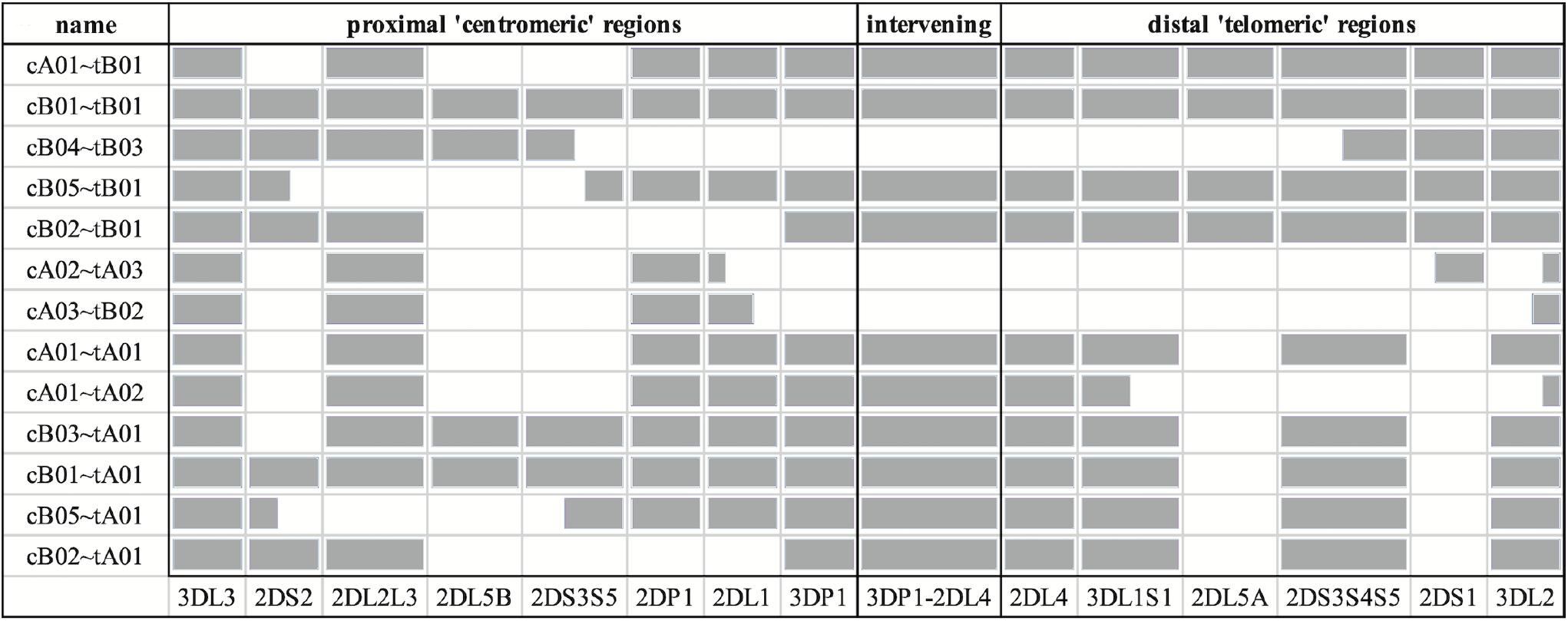
Schematic representation of the informal names and definitions of 13 human haplotype structures. The informal names of the haplotypes are in the first column. The definitions of the centromeric and telomeric regions are in the top row. The abbreviated names of the genes are in the bottom row; the intergenic region between *KIR3DP1* and *KIR2DL4* is represented as ‘3DP1-2DL4’. Grey cells indicates the presence of alleles for that gene; white indicates absence of an allele. Some cells are partially colored to indicate two-gene fusion alleles. The names of the haplotypes, as well as the gene content and locations are taken from the human curated annotations in the human genome reference. Haplotype cA01~tB04 contains a large insertion and was excluded for visualization.

### Workflow Overview

The result of the workflow was a multiple sequence alignment of 68 human and 1 chimp haplotypes. As shown in Figure 2, the first step was to align the 18 capture probes to the haplotype sequences (Figure 2A). Next, the sequence between each pair of probes (e.g., probe 4 to probe 3) was assigned a letter (e.g., C) (Figure 2B). In this way, each haplotype can be represented by a string (‘motifs’, e.g., ‘…CIJKL…’) whose length is 105 characters or less. These haplotype motifs were aligned to produce a high-level MSA (Figure 2C). The motifs allowed loci to be located in the haplotype sequences. The sequences for all alleles of each locus was aligned separately, and then the MSAs were concatenated to create the full haplotype DNA MSA (Figure 2D). In this two-alignment approach, the motif alignment provides structural annotation and the locus-specific DNA alignment provides base-level accuracy. The following paragraphs detail each step.

**Figure 2.**
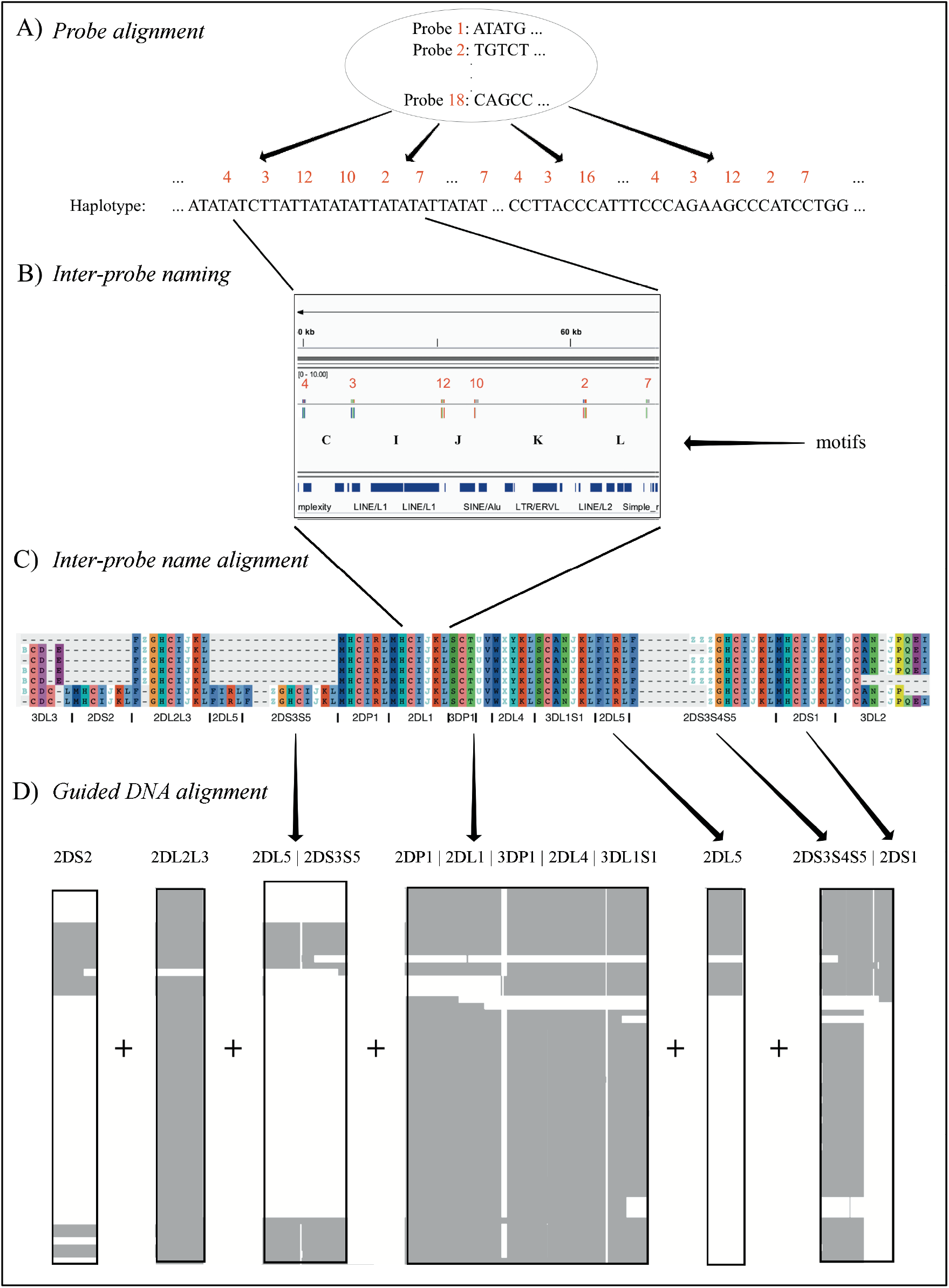
MSA workflow. Step **(A)** depicts the alignment of 18 120 base probes to each haplotype sequence. Step **(B)** shows how each ordered pair of probes in the alignment is assigned a letter. For example, the letter ‘I’ represents the sequence between the alignment of probe 3 and probe 12. Step **(C)** shows how each haplotype is depicted by a string of letters, how a haplotype motif MSA is generated from them, and how loci are defined in the MSA. Step **(D)** shows how the DNA in the motif-defined loci were separately aligned and then joined to create the final MSA.

### Probe alignment and inter-probe naming

The assembler preprint describes a method that uses 18 120 bp probes to capture KIR PacBio long-read sequences and assemble them into haplotypes with an average of 97% coverage and 99.7% concordance compared with reference sequences. When the probes are aligned to the haplotypes (Figure 2A), they align every 2398 bases on average. Probe locations are discovered by alignment with bowtie2 with options ‘-a --end-to-end --rdg 3,3 --rfg 3,3’. The alignment order of the 18 probes across a haplotype sequence allows that haplotype structure to be succinctly annotated as sequences of probe pairs (Figure 2B). For example, assume the alignment of probe 4 followed by probe 3 is called ‘C’, probe 3 followed by probe 12 is called ‘I’, and probe 12 followed by probe 10 is called ‘J’. Then the region ‘probe 4 to probe 3 to probe 12 to probe 10’ can be called ‘CIJ’. Probe 1 repeating once in the alignment is ‘Z’ and twice is ‘ZZ’, etc. There are 42 such probe pairs in this collection of 69 haplotypes. In this way, haplotypes can be briefly annotated as strings of a 42-character alphabet. See Figure 3 for an example of KP420440, whose full motif is MHCIJKLFGHCIJKLAIRLFZGHCIJKLMHCIRLMHCIJKLSCTUVWXYKLSCNOJKLAIRLFZG HCIJKLMHCIJKLFPCNOJQ. The motif pairs are defined in Supplementary Table 1; the probe sequences are defined in the assembly manuscript.

**Figure 3.**
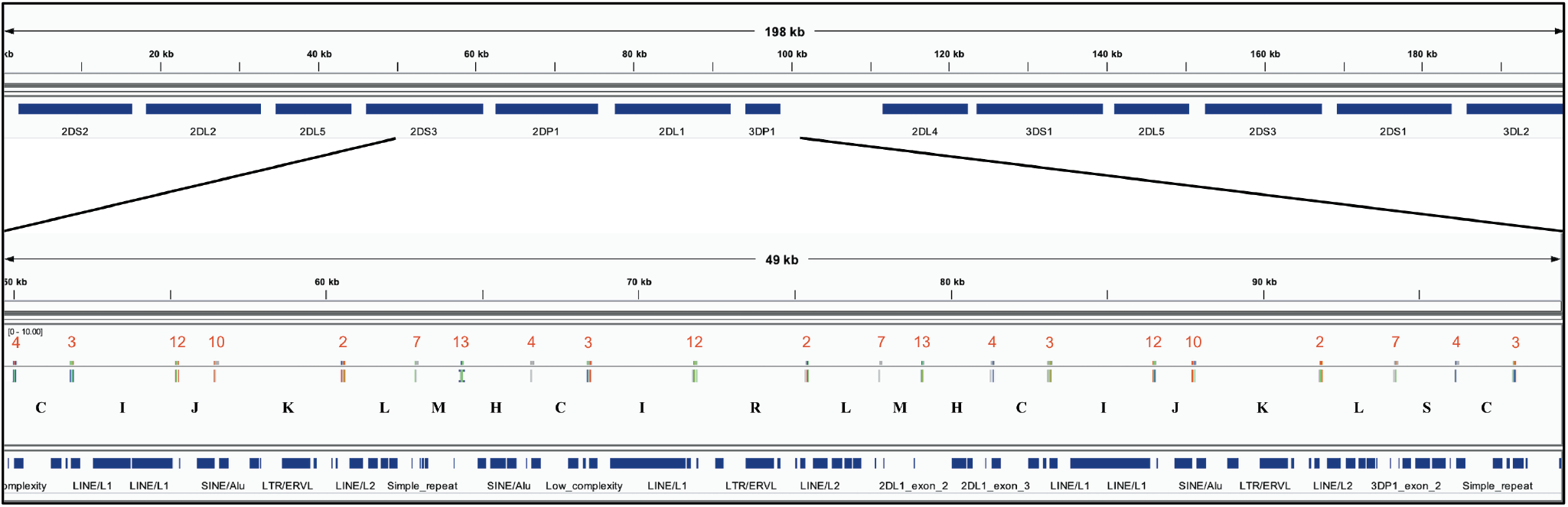
Example of the motif-generating alignment. The top frame shows the annotation for 198 kb haplotype KP420440 (cB01~tB01). The bottom frame zooms in to a 49 kb region depicting the alignment of the probes. In the middle of the bottom track, the locations of the probes are displayed by the vertical ticks with the red numbers above them and black letters below them. The red numbers label each probe, and the black letters label each probe pair (i.e., motif). The locations of exons and repeat elements (horizontal blue bars) are on the bottom track. Seven distinct probes align in this window. The motif pattern CIJKL occurs in two locations: 50-63kb and 81-94kb; the variation CIRL occurs 66-79kb.

### Inter-probe name alignment

A full-haplotype multiple sequence alignment of the probe motifs was generated for the 68 human haplotypes plus the chimpanzee as an outgroup (Figure 2C). Alignment of the motifs was created with MAFFT(10), but mostly aligned manually with Aliview 1.2.6; manual alignment was required to resolve ambiguities, as the motif alphabet is not directly supported by DNA or protein aligners. Ambiguous alignments were resolved by following the human curation of the reference haplotypes.

### DNA alignment

MAFFT was used to merge or add these haplotypes in the order cB02~tA01, cA01~tA01, cA02~tA01, cB02~tB01, cA0X~tB0X, cB0X~tA01, and cB0X~tB0X, where ‘X’ a general number not already used. Then, the full-length DNA sequences were separated into sets as defined by their motif structures (Figure 2D). Each set was aligned separately with MAFFT. The alignment was then edited manually by adding or deleting gaps to conform to locations in the probe motif MSA. Using the loci defined in the motifs, and the motif locations defined in the DNA, the alignment was refined for all alleles in each locus with MUSCLE in Aliview. A high-level depiction of the alignment was created with Jalview 2.11.0’s ‘Overview Window’ functionality and NCBI’s Multiple Sequence Alignment Viewer 1.13.1.

The MSA was validated at the structural level by showing that it recapitulates the human curation of the reference haplotypes in GenBank and the human genome reference. The alignment was validated at the allele level by showing each allele is assigned and annotated as expected with respect to IPD-KIR classifications.

The ability of existing software to align the 69 haplotypes was also evaluated with MAFFT stand-alone (--thread 19 --threadtb 10 --threadit 0 --reorder --adjustdirection --anysymbol -- leavegappyregion --kimura 1 ‒auto), PASTA v1.8.5 (docker run --rm -it -v $PWD:/opt/droe smirarab/pasta run_pasta.py), Clustal Omega v1.2.4 (clustalo ‒threads=29), M-Coffee metaserver 13.41.0.28, MUSCLE v3.8.31 (default parameters), and WebPrank (Updated 8 October, 2017). Implementation was conducted on the web servers when possible or a server running Ubuntu 18.04.4 LTS with 32 AMD Opteron(TM) Processor 6220 and 200 GB RAM and maximum java heap space set to -Xmx100G.

## Results

### Haplotype references

Haplotype sequences MN167504-MN167530 were deposited in GenBank; Supplementary Table 2 contains details. The 27 haplotypes include 8 cA01~tA01, 4 cA01~tA02, 4 cB01~tA01, 3 cB01~tB01, 3 cB03~tA01, 2 cA01~tB01, 2 cB02~tA01, and 1 cB01~tB01. Haplotype MN167506 (cB02~tB01), presumed to be of Asian ancestry, contains a *KIR2DL5B* allele (KIR2DL5B*00804) at the *KIR2DL5A* locus. Four haplotypes (MN167507, MN167509, MN167516, MN167530) with a *KIR3DL1*/*KIR3DL2* fusion(10) have been deposited in the human genome reference for the first time; they have been labeled as telomeric region ‘tA02’. Some of the diversity of structures in the African American cohort are depicted visually in Figure 4. In total, the 68 human haplotype structures consist of 27 cA01~tA01, 7 cB01~tA01, 6 cB01~tB01, 6 cB02~tA01, 5 cA01~tB01, 4 cA01~tA02, 4 cB03~tA01, 3 cB02~tB01, and 1 each for cA01~tB04, cA02~tA03, cA03~tB02, cB04~tB03, cB05~tA01, cB05~tB01.

**Figure 4.**
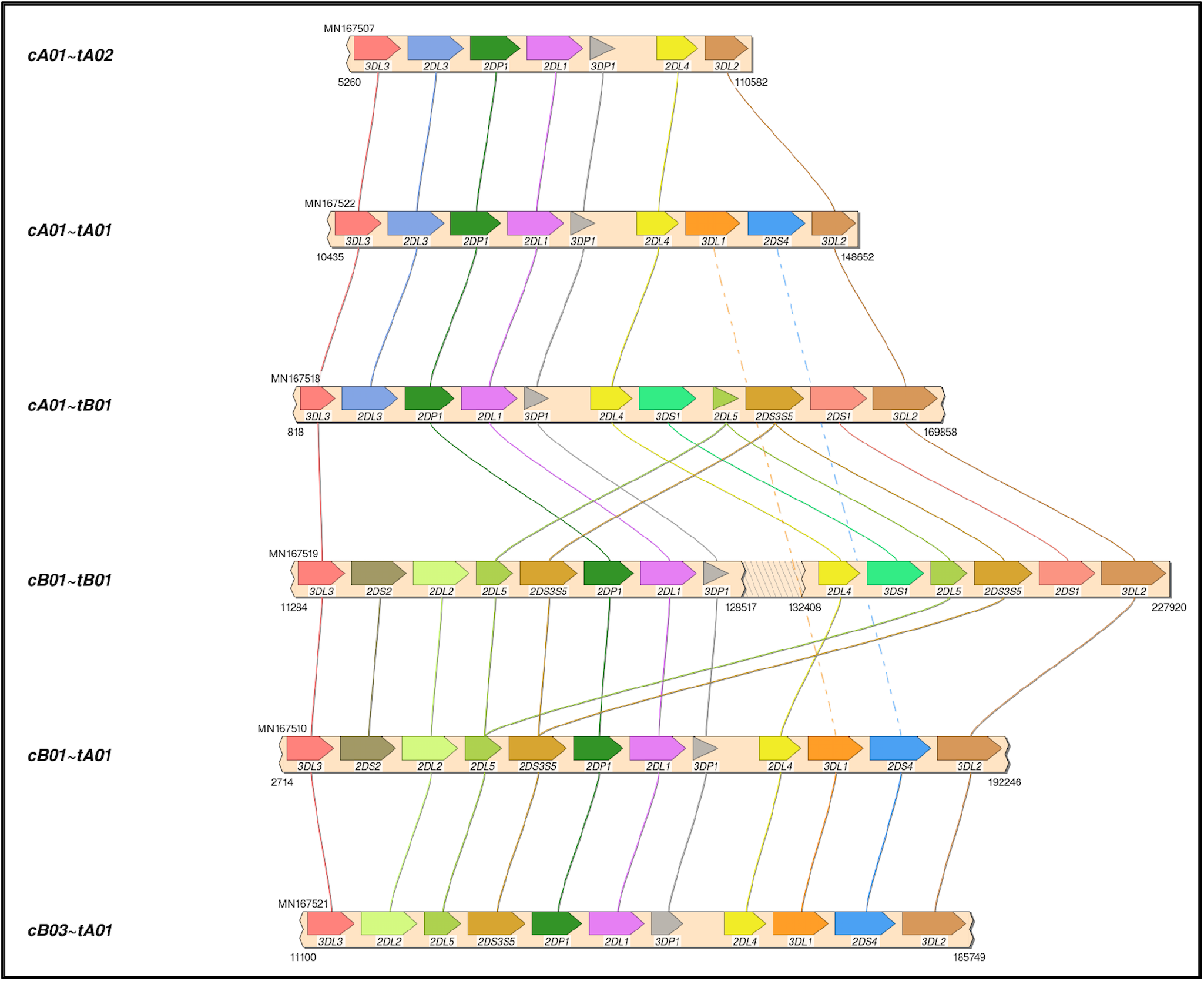
Haplotype structures in African American cohort. Shown are six of the eight structures in the African American cohort. Not shown are cB02~tA01, and 1 cB01~tB01.

### Validation of the MSA by annotation in human genome reference

Figure 5 shows a multiple sequence alignment of the probe motifs for the human haplotypes, excluding insertion-containing cA01~tB04 for space considerations. It recapitulates the human-annotated structures in Figure 1 with 105 positions. The common ~140 kb cA01~tA01 haplotypes are marked up in ~63 motif characters, and the ~220 kb cB01~tB01 haplotypes are marked up in ~94 motif characters. The average distance between probes is 2398; the maximum distance for non-*KIR3DL3* genes is ~5700 and for *KIR3DL3* it is ~7800. The haplotype motifs subdivide into 14 genic and 1 intergenic (*KIR3DP1*-*KIR2DL4*) loci. From 5’ (left) to 3’ (right) at the bottom of Figure 5, the 15 abbreviated loci are: 3DL3, 2DS2, 2DL2L3, 2DL5, 2DS3S5, 2DP1, 2DL1, 3DP1, 3DP1-2DL4, 2DL4, 3DL1S1, 2DL5, 2DS3S4S5, 2DS1, and 3DL2. The 14 genic loci consist of 9 distinct motifs: 2DL1, 2DS1, 2DS2 share MHCIJ; 2DL5A and 2DL5B share FIRL; 2DS3, 2DS4, and 2DS5 share FZ+GHCIJKL. The genes can be summarized as a short motif, or in some cases a regular expression. For example, FZ+GHCIJKL means: a F followed by one-or-more Zs followed by GHCIJKL. Supplementary Figure 1 contains the motif MSA from Figure 5. Supplementary Figure 2 adds cA01~tB04 (KU645196) and the chimpanzee haplotype to the MSA. The chimp haplotype contains 55 motif characters, encoding genes that align to the human *KIR3DL3~KIR2DS2~KIR2DP1~KIR2DL1~KIR3DP1~KIR2DL4~KIR3DL1S1~KIR3DL2*.

**Figure 5.**
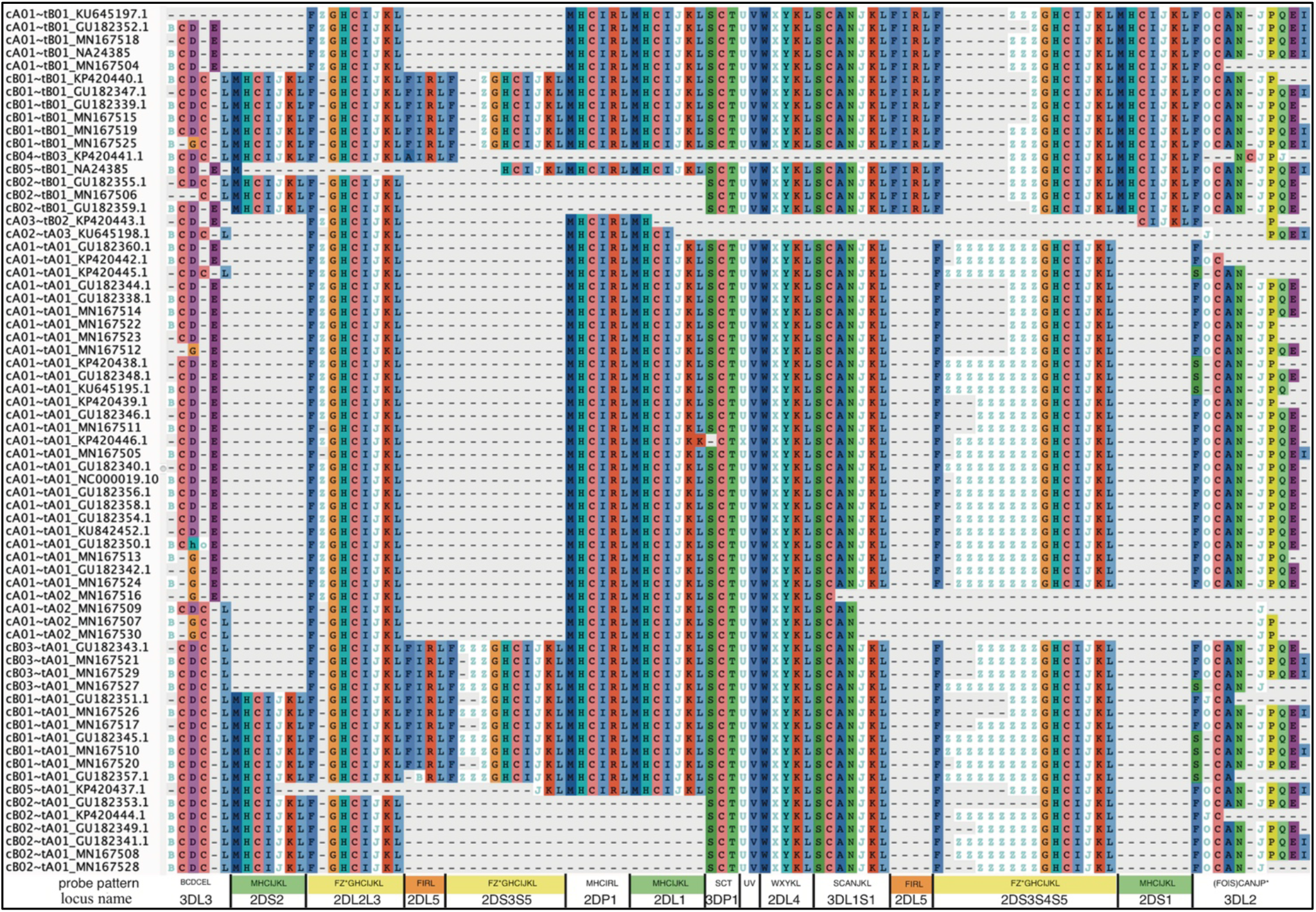
Probe motif multiple sequence alignment of the human haplotypes. The informal haplotype name and GenBank accession number are in the first column. The consensus motif pattern with each abbreviated locus name is described in the bottom row. Some loci share motifs: *KIR2DL1*, *KIR2DS1*, *KIR2DS2* (green); *KIR2DL5A* and *KIR2DL5B* (orange); *KIR2DS3*, *KIR2DS4*, and *KIR2DS5* (yellow). Some gene names are combined when they align at the same locus: *KIR2DL2* and *KIR2DL3* (2DL2L3), *KIR2DS3* and *KIR2DS5* centromeric side (2DS3S5), *KIR2DS*3, and *KIR2DS4*, and *KIR2DS5* telomeric side (2DS3S4S5), and *KIR3DL1* and *KIR3DS1* (3DL1S1). Haplotype cA01~tB04, which contains a 106 kb insertion, was omitted for visualization.

Haplotype sizes by number of loci (not including the *KIR3DP1*-*KIR2DL4* intergenic region) range from 4 loci (cA04~tA03) to 18 (cA01~tB04). Figure 6 displays a phylogenetic tree from the motifs of the haplotypes, including cA01~tB04 and the chimp haplotype; the aligned clusters recapitulate the haplotype categories in Figure 5.

**Figure 6.**
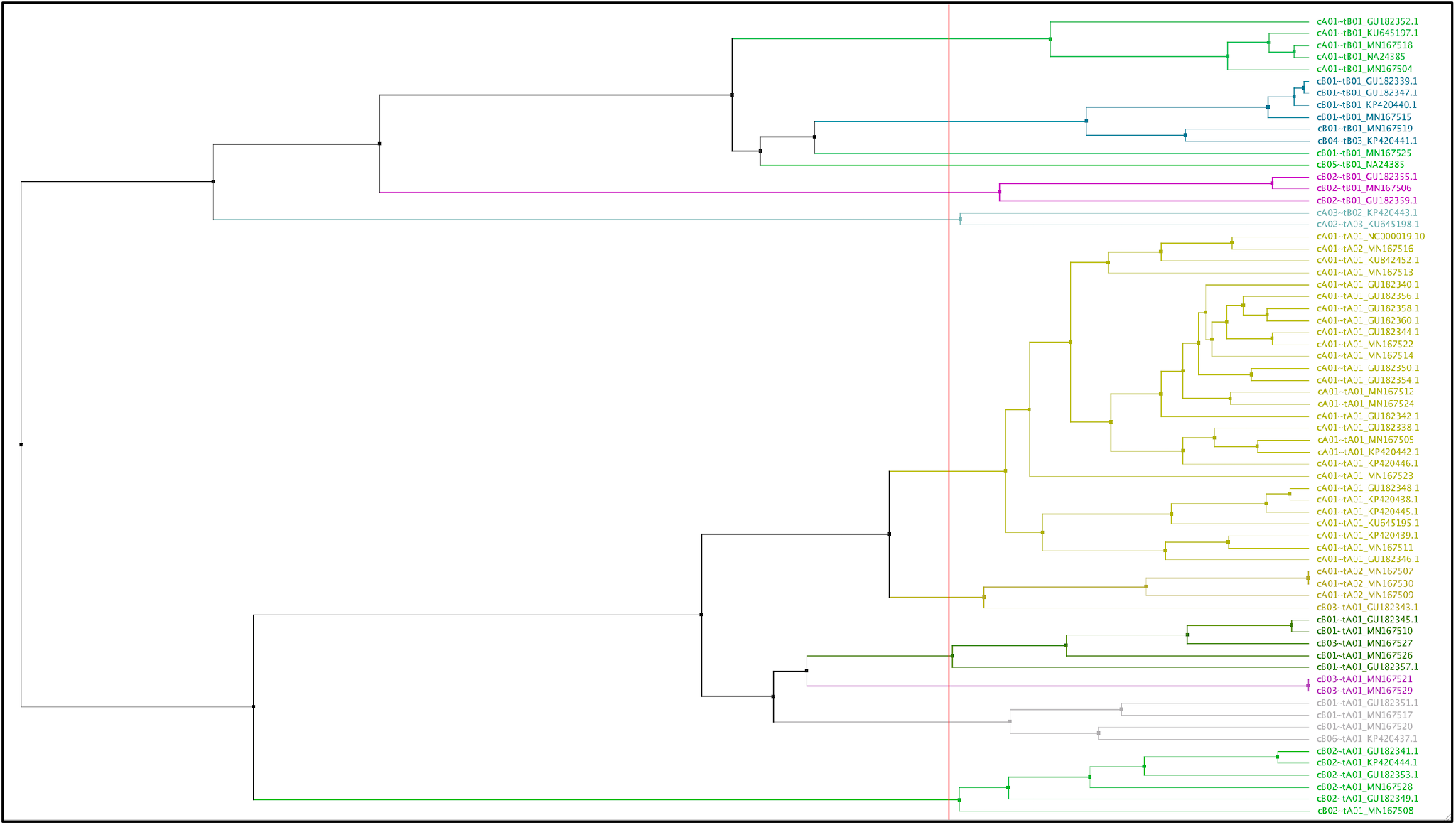
Phylogenetic tree made from the MSA from the motifs of 68 human haplotypes and 1 chimpanzee. The leaf descriptions on the right include the informal name of the structural haplotype as documented in GenBank as well as its accession numbers. The leaves are colored by their alignment order and the grouping defined by the red vertical line. The tree may not reflect the evolutionary history.

The DNA alignment of the 67 human haplotypes (minus cA01~tB04) has 263556 positions; it is included in Supplementary Figure 3. Figure 7B shows an overview of the DNA haplotype alignment as generated by Jalview’s ‘Overview Window’ function; the names of the haplotypes were added after exporting the overview image from Jalview. The pattern of white (gaps) and grey (aligned sequences) mirror the patterns in the schematic depictions of the human curated haplotypes in Figure 1 (recapitulated in Figure 7A). This evidence shows the DNA alignment matches the human curation at a structural level.

**Figure 7.**
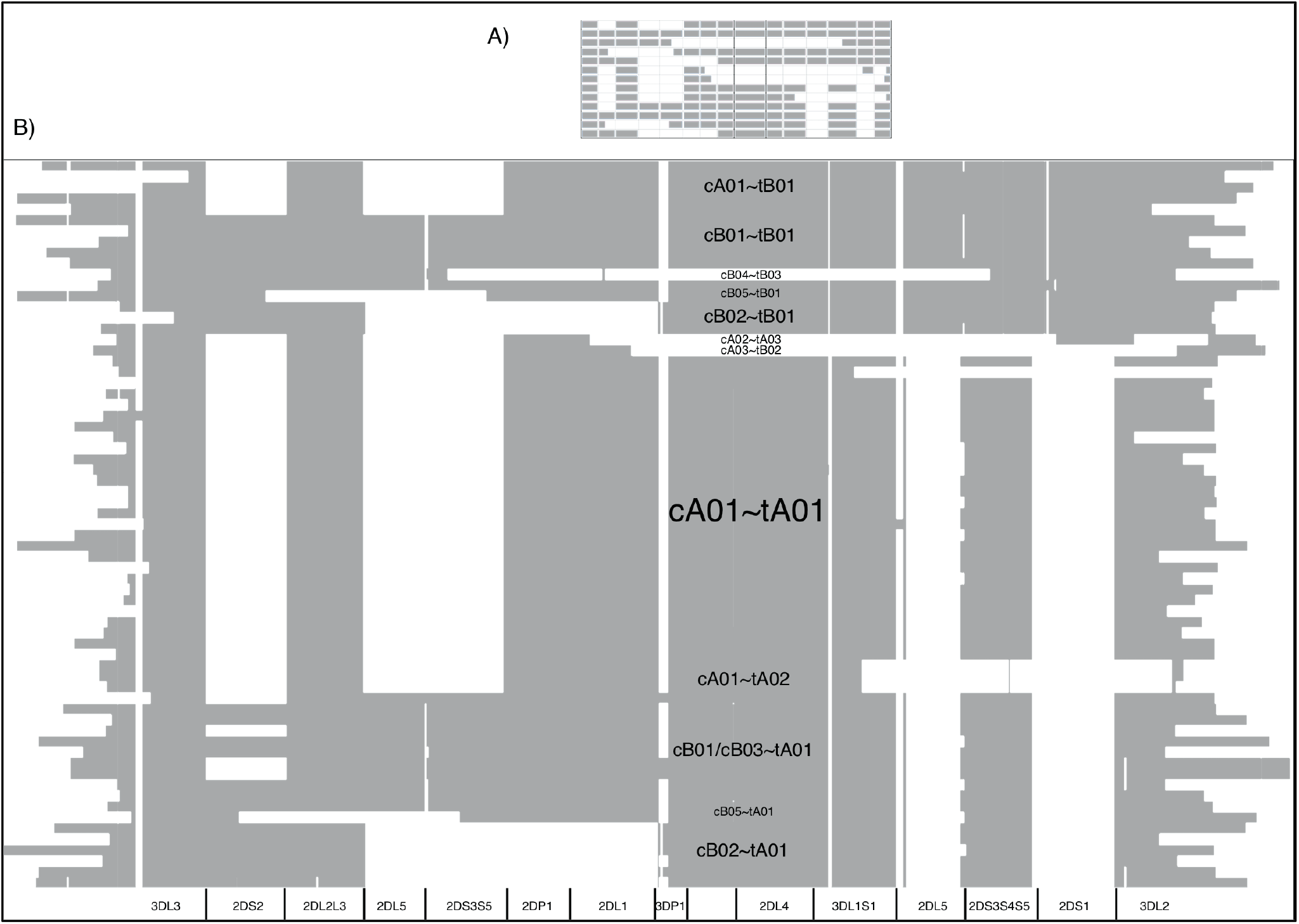
DNA MSA overview. The cartoon of the GenBank annotations from Figure 1 is in **(A)** and is included to compare with the Jalview overview of the DNA full-haplotype MSA from the 67 haplotypes in **(B).** The grey denotes alignment of the DNA bases. White denotes gaps between the sequences. cA01~tB04 is excluded for better visualization.

Figure 8 shows a view of the alignment of the 67 haplotypes using NCBI’s MSA Viewer. Positions that are in an agreement with consensus are colored in gray, positions that are not in an agreement with consensus are colored in red. In this tool, consensus includes gaps (lack of an allele at that position). For KIR, this causes the regions specific to B haplotypes to show as pure red, and the other regions to show as mixture of red and grey.

**Figure 8.**
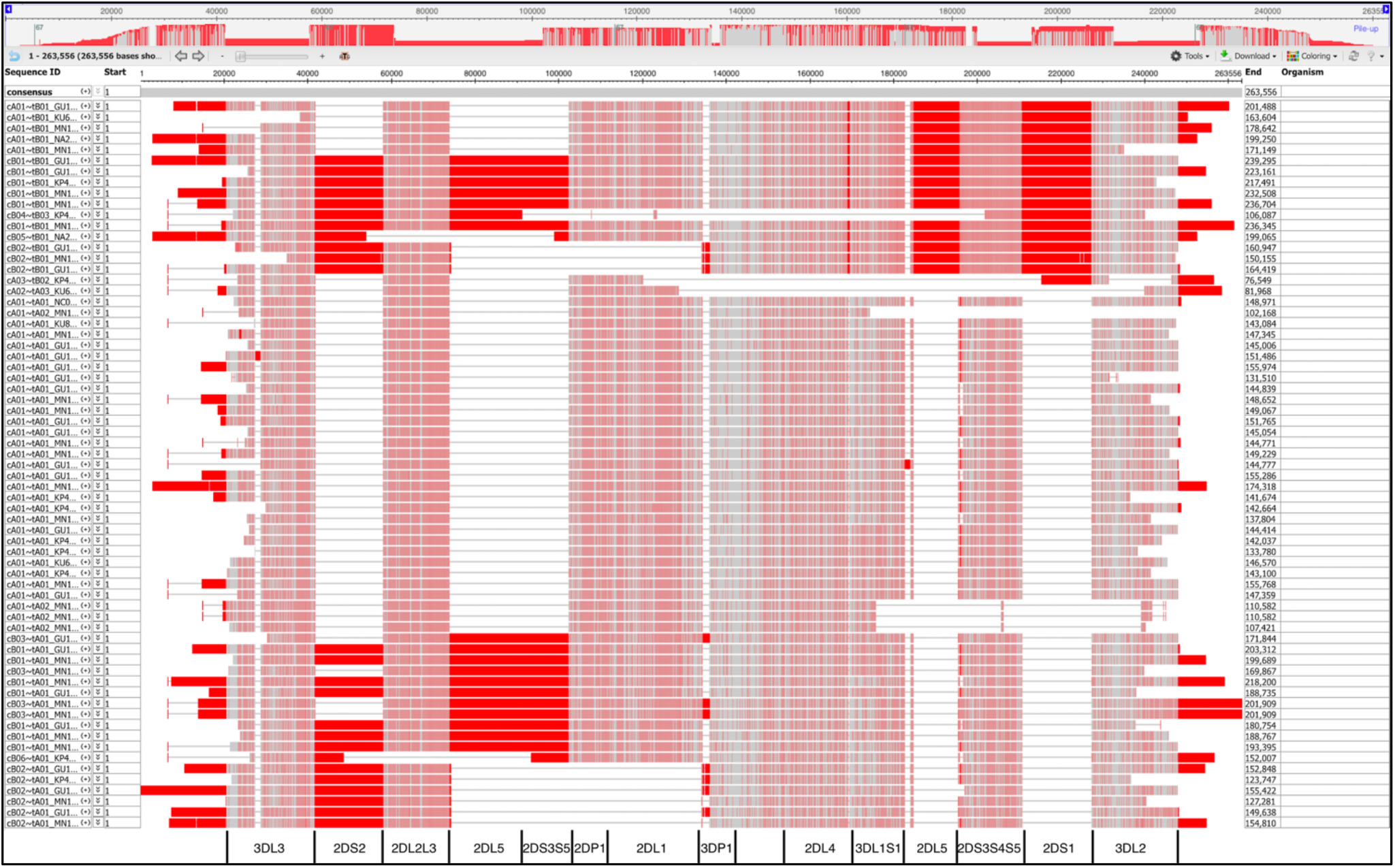
MSA Viewer depiction of the DNA alignment of 67 human haplotypes. Positions that are in an agreement with consensus are colored in gray, positions that are not in an agreement with consensus are colored in red. The location and borders of the genes have been added at the bottom.

### Validation of the MSA defined allele assignments by IPD-KIR annotation

The haplotypes are annotated by the names and orders of their alleles in Supplementary Table 2. Of the 647 alleles in the 68 haplotypes, 556 (86%) can be assigned names via IPD-KIR 2.9.0, at least at protein resolution. Unnamed alleles occur either because they have not yet been included in the IPD-KIR database, or they are partial alleles in the case of *KIR3DL3* and *KIR3DL2*. 97% of alleles can be named excluding *KIR3DL3*, *KIR2DP1*, and *KIR3DL2*. 39 *KIR2DP1* alleles are unnamed. To check the gene assignment for the 91 alleles that could not be named, the sequences were aligned to a set of 17 full gene alleles, each the first named allele for its assigned gene in IPD-KIR. Each allele being evaluated was considered to be correctly annotated if the allele sequence aligned closet to the IPD-KIR reference to which it was assigned by the motifs. Those results show that every unnamed allele aligned to the reference allele predicted by its motif assignment. The only exception was the GU182360 *KIR3DL2*, which aligns closest to *KIR3DP1*; however, this haplotype sequence is incomplete on the 3’ end, and the *KIR3DL2* allele only contains the 2221 5’-end sequences. Alleles that are a fusion of *KIR3DL1* and *KIR3DL2* (e.g. *KIR3DL1* alleles 059-061) are classified as *KIR3DL2* alleles using motifs but are classified as *KIR3DL1* alleles in IPD-KIR.

### Comparison with existing MSA software

Of the existing alignment software that was evaluated on the 67 human haplotypes, only stand-alone MAFFT was able to generate an alignment, using the FFT-NS-2 strategy. The input size was too large for Clustal Omega local server, M-Coffee web server, MUSCLE local server, and WebPrank web server, and the other algorithms ran out of memory. The MAFFT alignment, output, and overview are included in Supplementary Figure 4. Except for large portions of *KIR3DL3*, distal *KIR2DL3*/*KIR2DS4*/2DS3S5, and *KIR3DL2*, the MAFFT alignment does not recapitulate human-curated KIR haplotypes or gene alleles. It is 75% larger than our motif-guided alignment (960221 vs 263556 positions), and its overview shows the alignment columns are mostly gaps compared with the more expected block shaped column in Figure 7.

## Discussion

Although our evaluation of existing MSA software methods was not exhaustive, we believe it is unlikely that any current general-purpose alignment software can align all human KIR haplotype sequences consistent with the human curation. Alignment of the 68 sequences, each 67-269 kb, each homologous with itself and every other haplotype over lengths of 15 kb is very challenging without prior knowledge. Scoring matrices and algorithms like dynamic programming do not generally support sequences of this size under conditions where relatively young gene regions have duplicated both within and between haplotypes. To the best of our knowledge, a MSA of KIR haplotypes has never been published, despite the fact that dozens of haplotype sequences have been public for years.

Figures 5 and 6 demonstrate that the capture probe alignment motifs alone can annotate KIR haplotype sequences in a way that recreates the human curated annotation in GenBank. Figures 7 and 8 similarly show the accuracy of the DNA alignment; they demonstrate that the DNA annotated by the motifs recreates the assignment of the alleles in IPD-KIR. Also, for two practical examples, the capture probes have demonstrated the ability to assemble haplotypes with an average of 97% coverage (preprint), and the DNA MSA was used to discover gene markers that can genotype WGS with almost perfect accuracy(11) (preprint). The combination of multi-resolution validation and software applications demonstrates that both the motif and DNA alignments are likely to be accurate.

Figure 5 (alignment in Supplementary Figure 1) suggest 14 human KIR loci and 1 intergenic locus. The genic loci are abbreviated and labeled as 3DL3, 2DS2, 2DL2L3, 2DL5B, 2DS3S5, 2DP1, 2DL1, 3DP1, 2DL4, 3DL1S1, 2DS3S4S5, 2DS1, 3DL2, and the intergenic locus is between 3DP1 and 2DL4 (3DP1-2DL4). There are 9 motif-defined genes in the 14 genic loci (Figure 5). Loci abbreviated 2DL1, 2DS1, and 2DS2 share motif MHCIJKL. Loci 2DL2L3, 2DS3S5, and 2DS3S4S5 share variations of the FZ*GHCIJKL motif. Loci 2DL5A and 2DL5B share variations of FIRL. The motif MSA suggests *KIR2DS4* on the A haplotype shares a locus with *KIR2DS3* and *KIR2DS5* (not *KIR2DS1*) on the B haplotypes. The MSA similarly suggests *KIR2DL2* and *KIR2DL3* share a locus as do *KIR3DL1* and *KIR3DS1*. The Z character is unique to centromeric 2DS3S5 and telomeric 2DS3S4S5 loci and marks the alleles to a certain extent. In both, the *KIR2DS3* alleles have 1 Z in their motifs, *KIR2DS5* has 2 or 3, and *KIR2DS4* alleles are more variable.

Although the motifs correctly annotate the haplotype structures, they do not always do so unambiguously, nor do they always agree with the existing nomenclature as utilized in IPD-KIR. In the current gene nomenclature, there are 16 KIR genes. Some researchers have previously considered *KIR2DL2* and *KIR2DL3* to occupy the same locus as well as *KIR2DS3* and *KIR2DS5*. Conversely, *KIR2DL5* was once considered one gene but was split into *KIR2DL5A* for alleles in the centromeric locus and *KIR2DL5B* for alleles in the telomeric locus, creating new ‘A’ and ‘B’ designations that are independent from the ‘A’ and ‘B’ haplotype classifications. The motif MSA suggests the KIR region has 14 loci in 9 gene motifs. Under the existing gene nomenclature, *KIR3DL1* and *KIR3DL2* differ only by an index, since they are both three domain (“3D”) long tail (‘L’) genes; since the extracellular domains of *KIR3DL1/KIR3DL2* fusion alleles are from *KIR3DL1*, those fusion alleles are labelled in IPD-KIR as *KIR3DL1*. They are considered *KIR3DL2* alleles under the motif structure because the proximal portions of *KIR3DL1* and *KIR3DL2* share variations of SCANJ, but the distal portions are different; since the fusions are comprised of proximal *KIR3DL1* and distal *KIR3DL2*, they share a partial proximal motif with *KIR3DL1* and a full motif with *KIR3DL2*. They motif ambiguity between the *KIR3DL1/KIR3DL2* fusion and *KIR3DL2* can be resolved by linkage disequilibrium with *KIR2DS4*, since the fusion lacks *KIR2DS4* and the haplotypes with *KIR2DS4* cannot contain the fusion. Both systems consider the cB05 *KIR2DS2/KIR2DS3* fusion to be *KIR2DS2* (KIR2DS2*005 in IPD-KIR).

A total of 27 new haplotypes were added to the human genome reference as part of this study (accessions MN167504-30), 24 of which are from African Americans. The haplotypes cluster by structure, not population. Haplotypes from Africans or African Americans now constitute 47% of the KIR alternate references in the human genome project, and the human genome project contains more than three times as many alternative references for KIR than any full chromosome. These new haplotypes contain the first deposits of unusual linkages such as KIR2DL5B*00804 in the *KIR2DL5A* locus (MN167506), *KIR3DL1* or *KIR3DL2* (fusion) without *KIR2DS4* (tA02), *KIR2DL2* without *KIR2DS2* (cA03), and a *KIR2DS2* (fusion) without *KIR2DL2* (cB05). MN167526’s *KIR2DS2* allele has a 1 base deletion after 285th base in the CDS sequence, which leads to a premature stop code at position 435 in the CDS.

Scisco Genetics’ contribution to the human genome project of the KIR haplotypes has – like the human genome project itself – been an impressive feat on its own and will support the downstream discovery of the human immune system for years to come. Building on that work, the motifs and alignments presented here provide a means to help unify interpretation of the entire KIR region. They can be used to precisely define KIR haplotypes and loci, provide context for assigning alleles (especially fusion alleles) to genes, improve evolutionary inferences, improve imputation, interpret co-expression, and generate markers. The motif probes have been applied to a workflow to capture, assemble, and annotate KIR haplotypes at https://github.com/droeatumn/kass; it includes the ability to annotate KIR contigs/haplotypes as a separate workflow. The DNA alignments have been applied to discover markers used in a workflow to genotype KIR presence/absence from WGS at https://github.com/droeatumn/kpi.

## Author Contributions

DR and RK designed the experiments. DR performed the *in silico* experiments. CWP and DEG carried out KIR haplotype sequencing. All authors wrote the manuscript.

## Data Availability Statement

Haplotype sequences MN167504-MN167530 were deposited in GenBank. All haplotype sequences were referred to by their GenBank accession numbers, except the two haplotypes for isolate NA24385, which were provided as supplementary material the assembly manuscript preprint.

## Supplementary Table Captions

Supplementary Table 1. Motif assignments. The first column lists the alignment order of a pair of probes. The second column assigns a character to the pair.

Supplementary Table 2. Allelic haplotypes. Each row represents a haplotype and columns D-X contain the presence/absence of the gene and allele names from IPD-KIR 2.9.0. Column A contains and ID consisting of the haplotype structure and GenBank accession. Column B contains the population of the individual. Column C contains the GenBank accession of the haplotype.

